# Lysophosphatidic acid produced by Autotaxin acts as an allosteric modulator of its catalytic efficiency

**DOI:** 10.1101/346288

**Authors:** Fernando Salgado-Polo, Alex Fish, Minos-Timotheos Matsoukas, Tatjana Heidebrecht, Willem-Jan Keune, Anastassis Perrakis

## Abstract

Autotaxin is a secreted phosphodiesterase that converts lysophosphatidylcholine (LPC) into lysophosphatidic acid (LPA). LPA controls key cellular responses such as migration, proliferation and survival, implicating ATX-LPA signalling in various (patho)physiological processes and establishing it as a drug target. ATX structural and functional studies have revealed an orthosteric and an allosteric site, the “pocket” and the “tunnel”. Here, we revisit the kinetics of the ATX catalytic cycle in light of allosteric regulation, dissecting the different steps and pathways that lead to LPC hydrolysis. Consolidating all experimental kinetics data to a comprehensive catalytic model supported by molecular modelling simulations, suggests a positive feedback mechanism, regulated by the abundance of the LPA products activating hydrolysis of different LPC species. Our results complement and extend current understanding of ATX hydrolysis in light of the allosteric regulation by produced LPA species, and have implications for the design and application of orthosteric and allosteric ATX inhibitors.

## Introduction

Autotaxin (ATX or ENPP2) is a secreted glycoprotein and a unique member of the ectonucleotide pyrophosphatase / phosphodiesterase (ENPP) family^1^. It is the only ENPP family member with lysophospholipase D (lysoPLD) activity (*EC* 3.1.4.39), and it is the main enzyme responsible for the hydrolysis of lysophosphatidylcholine (2-acyl-*sn*-glycero-3-phosphocholine or LPC) to produce the bioactive lipid lysophosphatidic acid (monoacyl-sn-glycerol-3-phosphate or LPA) ^2–4^. LPA acts as a ligand for several LPA receptors (LPARs) showing overlapping activities. The ATX-LPA signalling axis is vital for embryonic development and has been implicated in many (patho)physiological processes, which include vascular development^5^, cancer metastasis^6^, and other human diseases, such as fibrosis^7^ and cholestatic pruritus^8^.

ATX is translated as a pre-proenzyme that is secreted to plasma upon its proteolytic processing, resulting in its native structural domains ^9,10^. Close to the N-terminus, ATX presents two somatomedin B (SMB)-like domains, which are followed by the central catalytic phosphodiesterase (PDE) domain, and an inactive nuclease-like domain. Catalysis occurs in a bimetallic active site presenting two Zn^2+^ atoms, and resembles that of other members of the alkaline phosphatase family ^11^. The catalytic site of ATX is organized in a tripartite binding site (Fig.1), where the active site is followed by a shallow hydrophilic groove that accommodates the glycerol moiety of lipid substrates, and nucleotide substrates. This groove leads to a T-junction and two separate paths: a hydrophobic pocket where the acyl chain of the lipid substrate can bind ^11–13^, and a tunnel, also called in literature “hydrophobic channel”, leading to the other side of the PDE domain ^13^. The open tunnel is defined by the PDE and the SMB-1 domains and presents both hydrophobic and hydrophilic residues on its inner walls ^11^.

**Figure 1.**
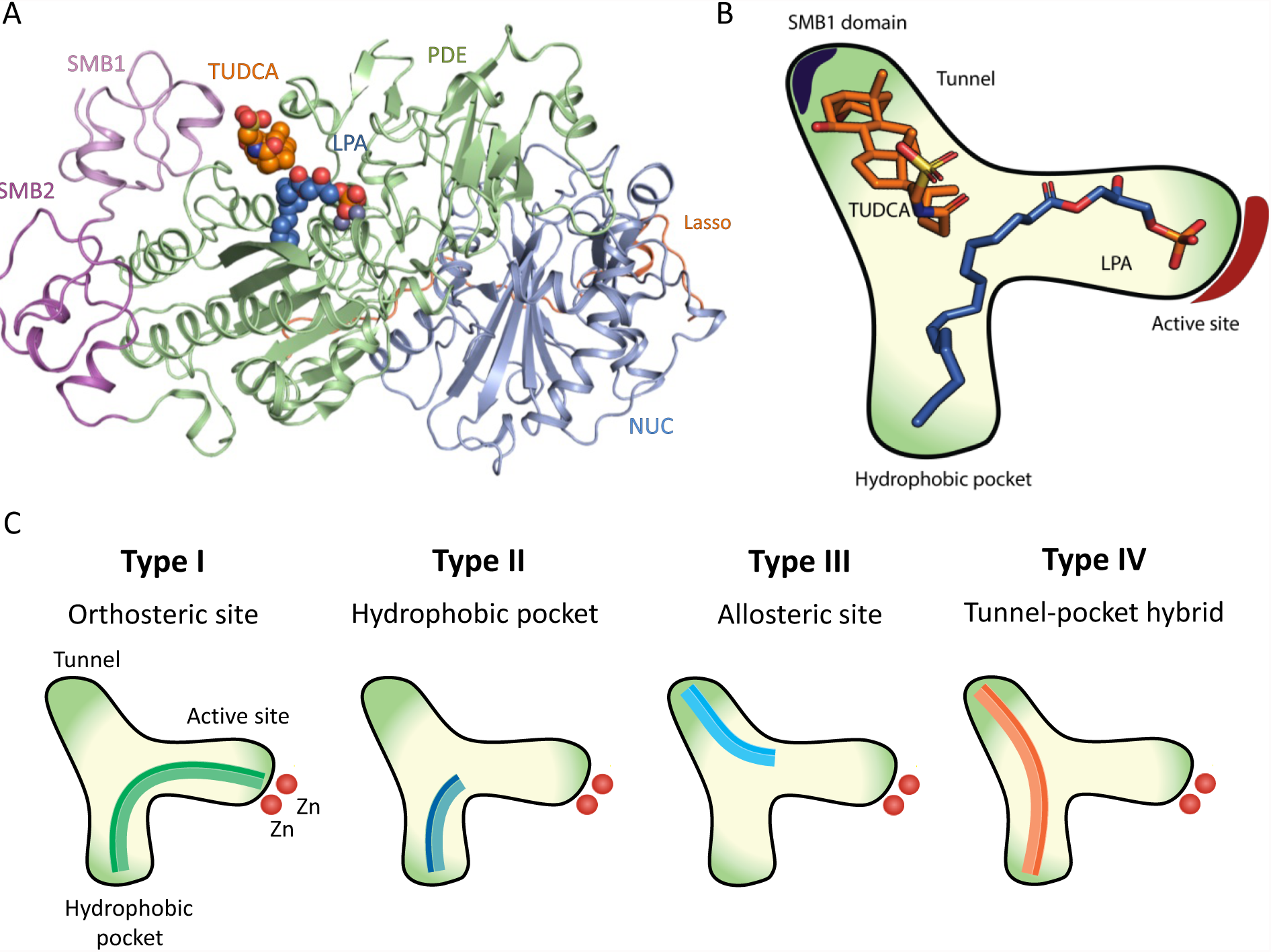
Structure of ATX and its tripartite site bound to LPA and the bile salt TUDCA. A, Crystal structure of rat ATX bound to bile salts (PDB ID 5dlw). ATX presents four distinct domains: two somatomedin B-like domains (SMB-1 and SMB-2), the active catalytic phosphodiesterase (PDE) domain, and an inactive nuclease-like domain. Additionally, the site where furin cleaves pro-ATX is indicated. B, cartoon representation of ATX tripartite site, consisting of a catalytic site, a hydrophobic pocket and a tunnel that acts as an allosteric regulator of ATX activity. C. The four types of ATX small molecule inhibitors: Type I inhibitors mimic the LPC mode of binding and include HA155 (IC_50_ = 5.7 nM) ^3^ and PF-8380 (IC_50_ = 1.7 nM) ^14^ which have been also validated to lower LPA levels in vivo. Type II ATX inhibitors obstruct lipid binding to the hydrophobic pocket, just like PAT-078 and PAT-494 ^15^. Type III inhibitors occupy the allosteric regulatory tunnel, modulating ATX activity by non-competitive inhibition, such as the steroids 7α-hydroxycholesterol, tauroursodeoxycholic acid (TUDCA) (IC_50_ = 11 µM) or ursodeoxycholic acid (UDCA) (IC_50_ = 9 µM) ^16^. Based on the binding mode of pocket-binding type II and tunnel-binding type III inhibitors, type IV compounds have been produced, either by design, fusing parts of a type II and a type III inhibitor, such as in compound 17 (IC_50_ = 14 nM) ^17^, or by serendipity followed by specific structure-based design, such as in GLPG1690^18^.

The structural characterization of the ATX tripartite site (Fig. 1A, B) has created a remarkable potential for selective inhibitor design ^15,19,20^ that includes lipid-based inhibitors ^21^, DNA aptamers ^22^ and small molecules. The latter can be classified in four distinct types depending on their binding modes to the tripartite site ^23^ (Fig. 1C). The work on type III and type IV inhibitors has highlighted further the tunnel as a novel allosteric site to modulate ATX activity. This modulation has been hypothesized to occur by disturbing the catalytic cleavage of substrates and/or by modulating LPA product uptake and the delivery to its cognate LPA receptors ^16^. In such a mechanism, ATX not only drives the formation of LPA but also ensures specificity in LPA signalling. However, the mechanisms underlying tunnel modulation have not been characterized.

ATX activity measurements often utilize specific synthetic unnatural fluorescent substrates that allow direct measurement of the phosphodiesterase activity ^24^: bis(*p*-nitrophenyl) phosphate ^25^, CPF4 ^26^, thymidine 5’monophosphate *p*-nitrophenyl (*p*NP-TMP) ^2^, the fluorogenic substrate 3 (FS-3) ^27^, and the fluorescent probe 12-(*N*-methyl-*N*-(7-nitrobenz-2-oxa-1,3-diazol-4-yl)) (NBD)-LPC ^28^. Albeit such synthetic substrates are invaluable for inhibitor development, kinetic studies that aim to describe the physiological activity of ATX, should focus on the natural substrates ^25,29^. Hydrolysis of the physiological substrate is commonly detected by a fluorimetric method consisting of two coupling enzymes: choline oxidase and horseradish peroxidase (HRP) (Fig. 2) ^30^. In this assay, choline oxidase converts choline into betaine and hydrogen peroxide, which HRP in turn uses to oxidize homovanillic acid (HVA), causing dimerization of HVA, allowing to reach its fluorophoric state (Scheme 1) ^30^. This indirect fluorescence-based method is the only assay to measure ATX catalytic activity towards its physiological substrate.

**Figure 2.**
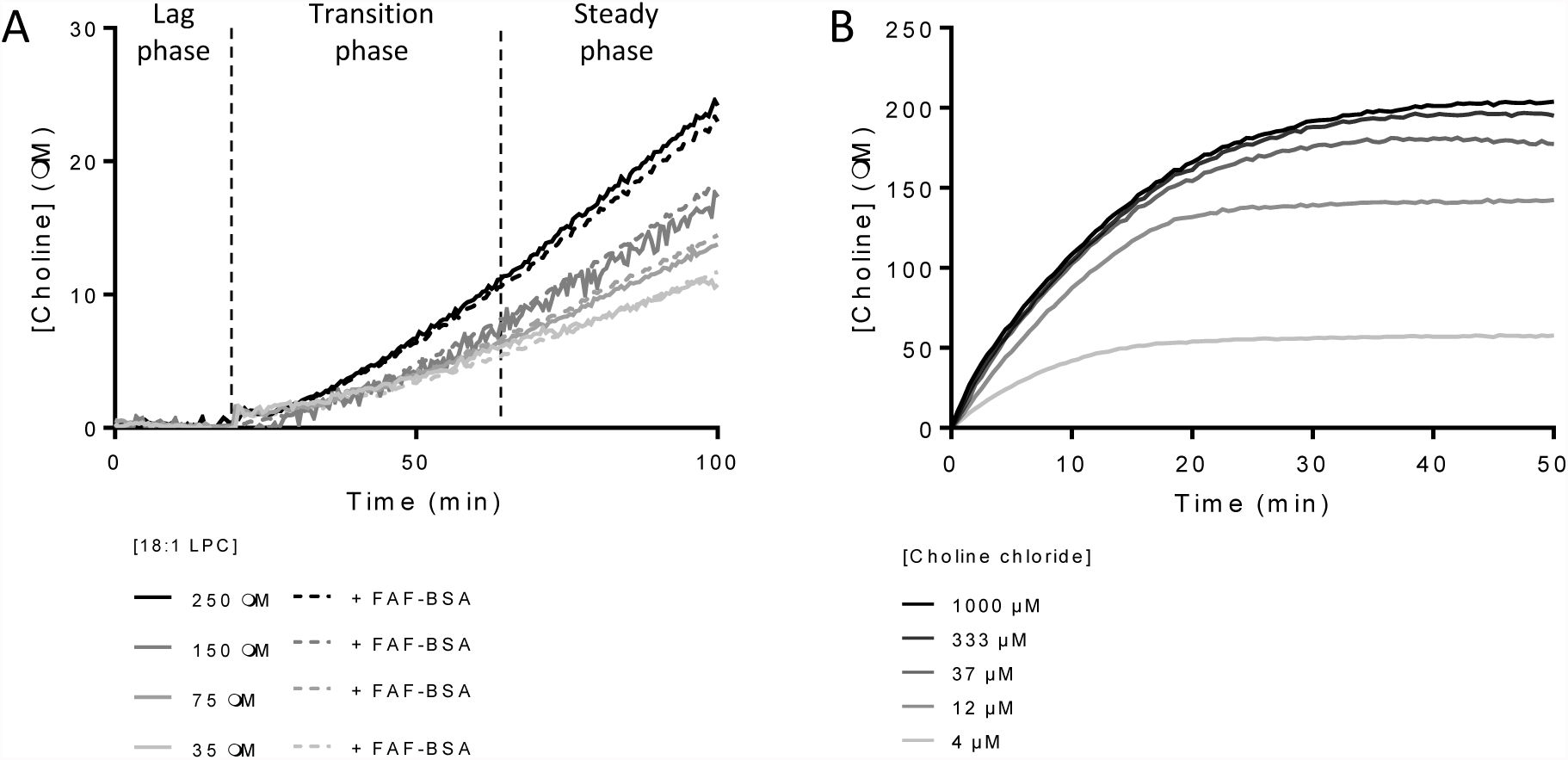
Enzyme kinetics described by ATX and choline oxidase for the hydrolysis of their physiological substrates. A. The kinetics for ATX lysoPLD activity on LPC results in a double exponential-sigmoid choline secretion that can be measured by a coupled reaction with choline oxidase and HRP. This behaviour is independent of the presence of delipidated BSA acting as an LPC vehicle B, Choline chloride oxidation by choline oxidase follows a single exponential kinetics. Data represent the averaged data of triplicate measurements.

**Scheme 1.**
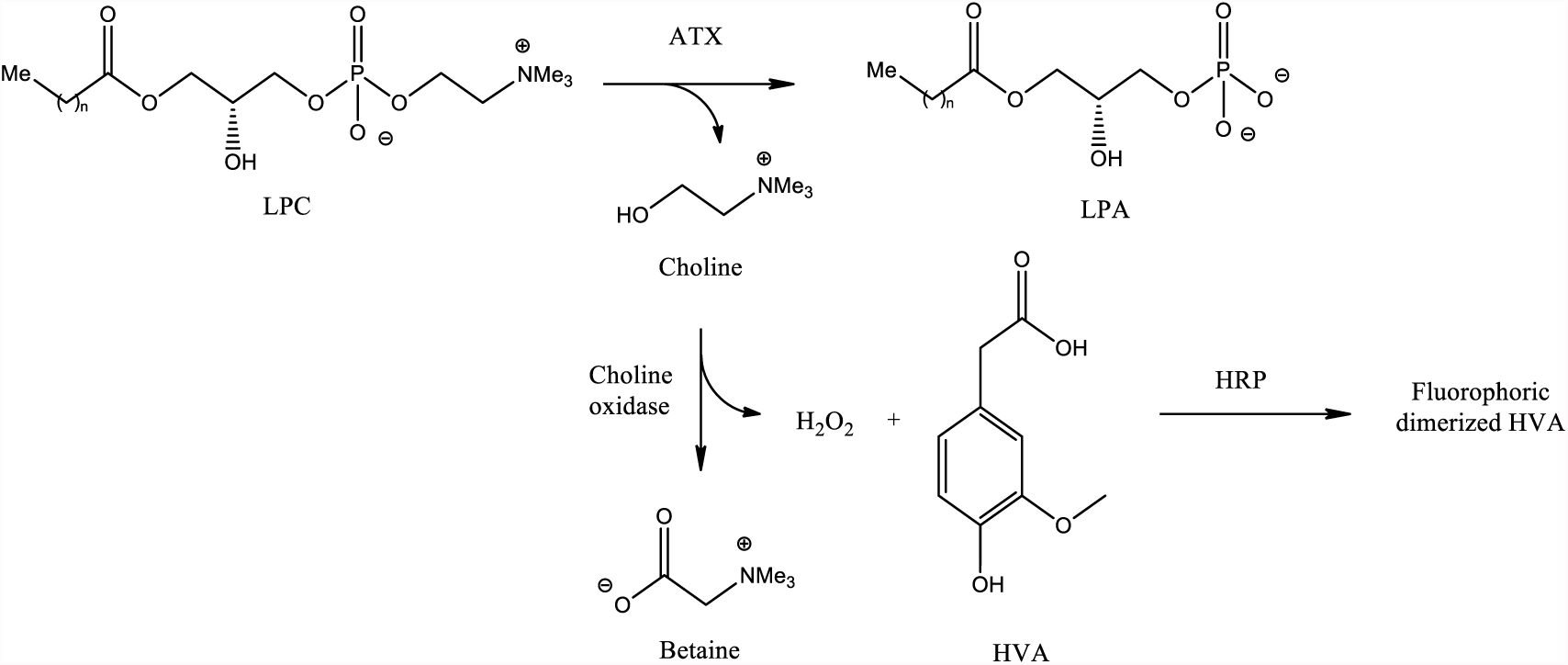
Choline release coupled enzymatic assay for measuring ATX lysoPLD activity. LPC, lysophosphatidylcholine; LPA, lysophosphatidic acid; HVA, homovanillic acid; HRP, horseradish peroxidase.

An early study on ATX activity had put forward a model where LPA acts as an inhibitor of catalysis ^25^. Albeit that stands unquestioned for artificial ATX substrates (e.g. *p*NP-TMP and FS3), recent experiments from other groups ^25,28^, could not corroborate a role of LPA in product inhibition of LPC hydrolysis. Importantly, a kinetic analysis by Saunders and colleagues reported on the substrate-specific kinetics of ATX towards NBD-LPC and FS-3 and provided valuable insight into the catalytic cycle of ATX ^28^. Their results showed that 12:0 NBD-LPC binding and hydrolysis were slow and rate-limiting, and provided clear evidence for a model where catalysis first leads to product release of choline and is followed by release of 12:0 NBD-LPA. This mechanism is also supported by subsequent structural studies that support an associative two-step in-line displacement mechanism ^19^. The slow release of LPA, taking place in the range of tens of seconds ^28^, suggested a mechanism where ATX could also act as an LPA carrier, spreading LPA signal to distal locations from those where LPC was taken. Such a model is also supported by data that show that Autotaxin can be recruited to the cell surface binding both to integrins ^13,31^ and surface heparin sulfate proteoglycans ^32^.

In this study, we set out to examine key questions about ATX catalysis that help explain the allosteric modulation of its physiological activity as a lysophospholipase D. We first establish that ATX activity is stimulated by LPA using the physiological LPC substrate and a new fluorescent substrate (NBD-LPE). Following a bottom-up approach, we then model and validate the lysoPLD activity of ATX on LPC. Next, we used ATX inhibitors from different classes and molecular dynamics simulations to propose that the LPA activates ATX lysoPLD activity by binding to the ATX tunnel. Our results propose a novel concept in the way we understand ATX hydrolysis, helping explain ATX function *in vivo*, and aiding future inhibitor design and optimization.

## Results

### ATX hydrolysis presents a lag phase

ATX lysoPLD kinetics on LPC, always present an initial lag phase, which is followed by a linear phase in which ATX reaches its maximal catalytic activity as LPC is consumed (Fig.2A). The recorded choline oxidation signal can be better described by a double exponential rather than a single exponential (P < 0.0001) that would be characteristic of Michaelis-Menten kinetics. As it is not clear whether this is an artefact of the coupled assay or some intrinsic LPA-mediated regulation of ATX catalytic cycle takes place ^30^, we set out to revisit the model for ATX catalysis.

We have first excluded trivial parameters such as changes in temperature or slight variations in buffer composition between different substrate concentrations, by carefully controlling experimental conditions. Importantly, the lag phase is observed in the absence or the presence of fatty acid-free (FAF) bovine serum albumin (BSA), implying that the initial lag phase is independent of the presence of a lipid vehicle. Then, we wanted to exclude the possibility that the observed behaviour of choline release, is due to the kinetics of oxidation of choline by choline oxidase (EC 1.1.3.17) yielding betaine and two H_2_O_2_ molecules (Scheme 1). Choline oxidation and subsequent HVA dimerization over time are described by hyperbolic kinetics (Fig. 2B) compatible with the Michaelis-Menten approximation. The parameters derived for this reaction (*K_M_* = [98 - 144 µM] and *kcat* = [1.28 - 1.36 s^-1^], Table 1) fit well to the ones reported for the enzymatic activity of choline oxidase in literature (*K_M_* =[0.1- 2 mM] and *k_cat_* [1- 3 s^-1^]) ^33^.

**Table 1.** Comparative table of the Michaelis-Menten-derived kinetic parameters of choline oxidation, and 18:1 LPC and 18:1 NBD-PEG4 LPE cleavage with respect to those reported for 12:0 lyso-NBD-PC^28^.

| Kinetic Parameters | Choline Oxidase | 18:1 LPC | NBD-PEG4 LPE | NBD-LPC |
| --- | --- | --- | --- | --- |
| $K_M (\mu\text{M})$ | $121 \pm 23$ | $112 \pm 22$ | $0.88 \pm 0.09$ | $308 \pm 195$ |
| $k_{cat} (\text{s}^{-1})$ | $1.32 \pm 0.04$ | $0.016 \pm 0.01$ | $0.021 \pm 0.003$ | $0.06 \pm 0.01$ |
| $k_{cat}/K_M (\mu\text{M}^{-1} \text{s}^{-1})$ | $1.08 \cdot 10^{-2} \pm 1.0 \cdot 10^{-4}$ | $2.1 \cdot 10^{-4} \pm 1.0 \cdot 10^{-5}$ | $2.38 \cdot 10^{-2} \pm 2 \cdot 10^{-4}$ | $1.9 \cdot 10^{-4} \pm 7 \cdot 10^{-5}$ |

**Table 2.**
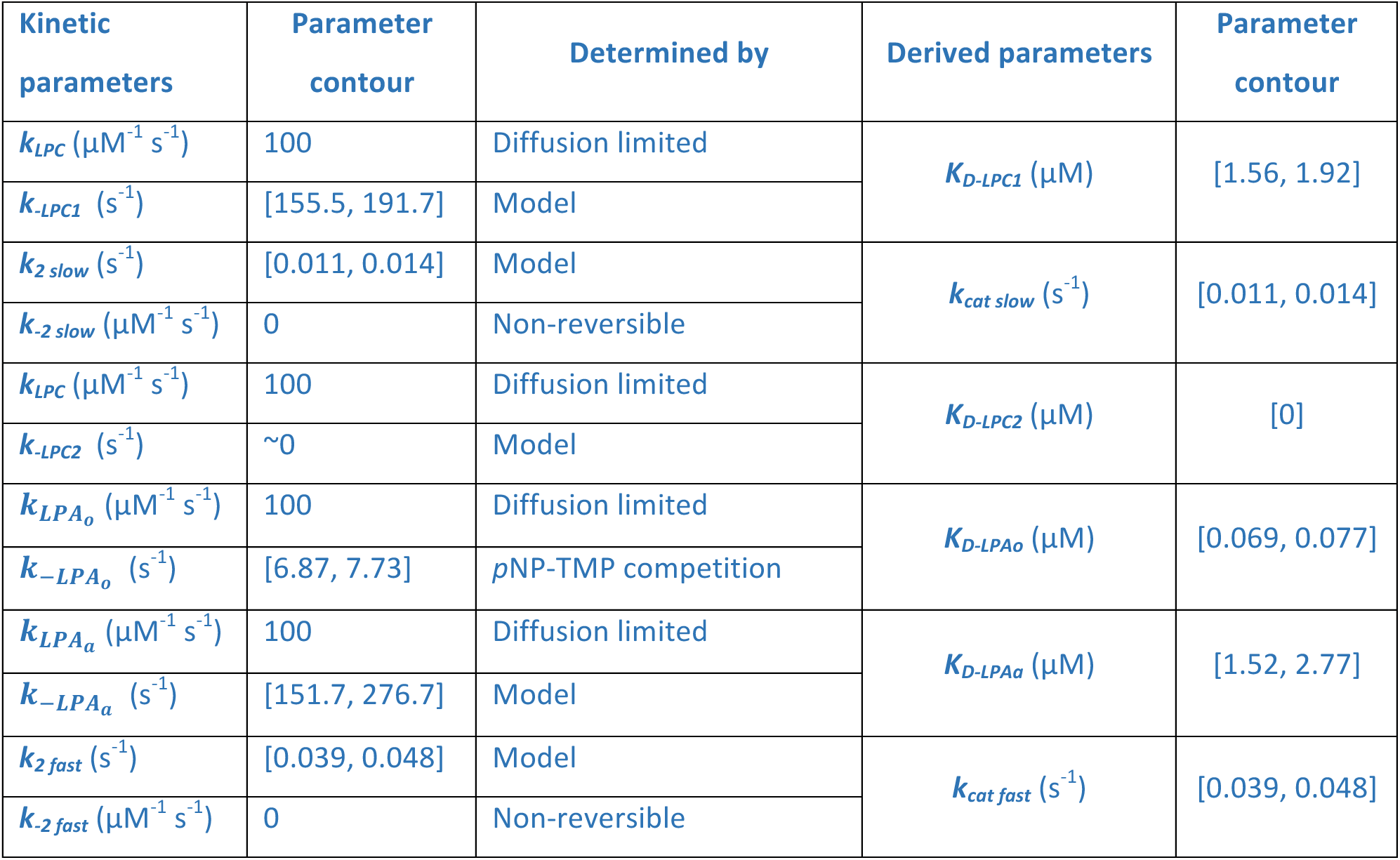
Kinetic parameters of the KinTek model in Figure 7. The confidence contours of each parameter were calculated employing FitSpace statistical analysis on Kin Tek at a chi^2^ > 0.9901 (supplemental figure 8).

To confirm the observed lag phase, we also wanted to use a direct readout assay using fluorescently labelled LPC. Existing fluorescently labelled substrates, such as 12:0 NBD-LPC, carry the label in the acyl chain, and are reporters of LPA binding and release ^28^ and thus not useful to measure the actual kinetics of catalysis and choline release in the first reaction step of the double displacement mechanism. To have a direct readout for choline release, Avanti Polar Lipids produced a new fluorescent substrate, namely 18:1 NBD-PEG4 LPE (Fig. 3A). This chromophoric substrate presents a PEG4 linker between the ethanolamine moiety and the fluorophore, which provided the necessary flexibility to be cleaved off by ATX (Fig.3B), which is promiscuous to the nature of the head group. Interestingly, the direct measurement of head group release revealed that the lag phase in the beginning of ATX catalysis was still present (Fig.3C). The linear phase of the experimental data was fitted by the Michaelis-Menten derivation to obtain the kinetic parameters for the cleavage of this substrate, indicating that the NBDPEG4-LPE *K_M_* is 100-fold lower than that of LPC, whereas *k_cat_* is in the same range. PF-8380 ^14^ and compound 17^17^ both inhibited NBD-PEG4-Ethanolamine release similarly to the choline release coupled reaction (Supplemental table 1), suggesting that the binding mode of NBD-PEG4-LPE is similar to LPC. While this substrate confirmed that the lag phase is a genuine feature of ATX-mediated LPC hydrolysis, it presented some solubility issues and a quenching effect at concentrations higher than 15 µM. Thus, we decided to use the native LPC substrate for further characterization.

**Figure 3.**
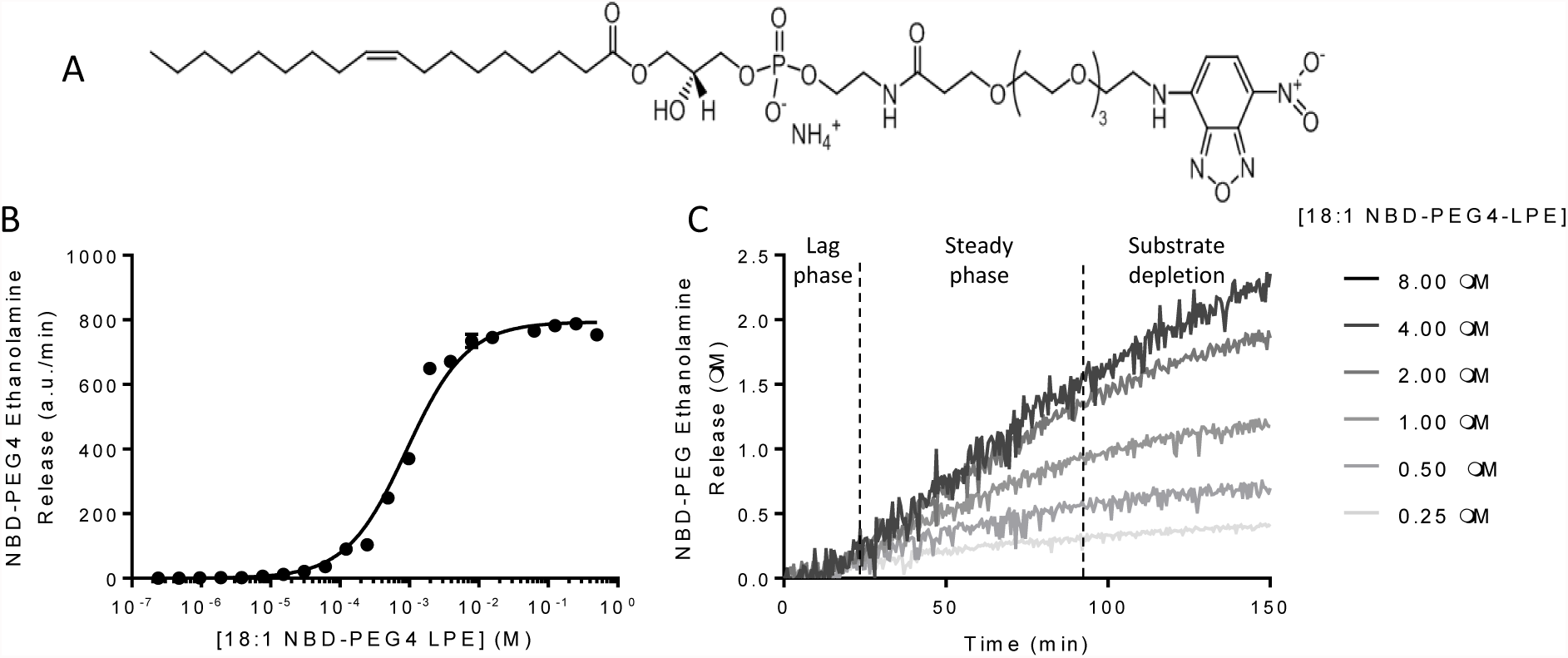
Cleavage of the novel probe 18:1 NBD-PEG4 LPE substrate by ATX lysoPLD activity presents initial lag phase. A, molecular structure of NBD-PEG4 LPE, presenting an optimal λ_exc_/λ_em_ = 468/546 nm. Product release was detected as a loss in fluorescence intensity at 546 nm with respect to that of the NBD-PEG4 LPE substrate. B, Substrate concentration curve fit to the Michaelis-Menten equation employed to determine the kinetic parameters of NBD-PEG4-LPE by measure NBD-PEG4 ethanolamine release. C, NBD-PEG4 ethanolamine release kinetics upon ATX lysoPLD activity, which follows a three-phase behaviour similar to that observed in Fig.2A. In all cases, data represents means of triplicates ± SEM.

### LPA modulates ATX catalysis

Having established the lag phase and subsequent activation of ATX-mediated catalysis, we wanted to examine a possible role of LPA in this behaviour. We first revisited the role of LPA in modulating hydrolysis of various ATX substrates, by adding increasing amounts of LPA in different reporter reactions indicative of ATX phosphodiesterase activity. Consistently with previous results ^13,25^, we observed that 18:1 LPA inhibited hydrolysis of bis-*p*NPP, CPF4 and *p*NP-TMP with an IC_50_ of ^~^50 nm (Fig.4A-B). This reduction in total ATX phosphodiesterase activity can be attributed to the competition exerted by LPA on active site-binding nucleotide substrates. However, analysing experimentally the cleavage of the physiological substrate 18:1 LPC while titrating increasing amounts of the homologous LPA species, we observed an activation of LPC hydrolysis up to ^~^50%. The half maximum activation was observed in an LPA concentration that we define as AC_50_, which was 1.4 ± 0.4 µM (Fig.4C). LPA modulation of ATX lysoPLD activity was also detected in the new direct readout assay. As observed in the choline release assay, activation by 18:1 LPA resulted in an AC_50_ value of 1.1 ± 0.3 µM (Fig.4D). Consistently, pre-incubating ATX with LPA for 30 minutes before supplying the LPC substrate reduced the length of the lag phase (Fig.4E-F).

**Figure 4.**
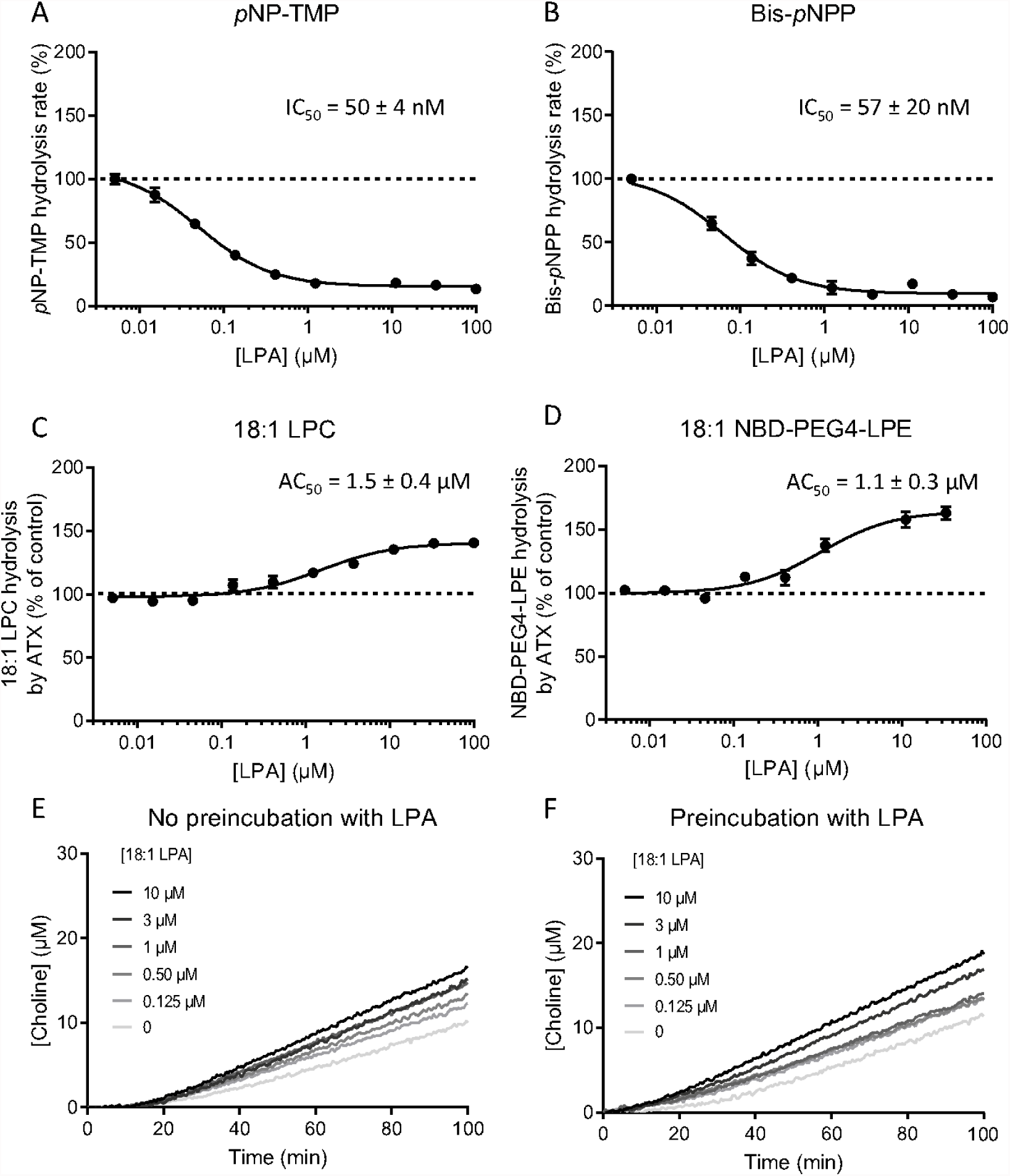
LPA modulates ATX catalysis, where it can act as an activator or inhibitor in a substrate-specific manner. A, B, LPA inhibition of the hydrolysis of the unnatural ATX substrates pNP-TMP and bis-pNPP. IC_50_ values were obtained from these fits using Equation 5. C, effect of the addition of 18:1 LPA on human ATXβ lysoPLD activity. LPA induced up to 45% increase in activity with respect to the control reaction. D, activation of the lysoPLD activity of ATX activity by 18:1 LPA using 18:1 NBD-PEG4 LPE as substrate yielded the same activation constant. This result confirmed that data obtained with the coupled assay is reproducible with the novel substrate. E, F, The lag phase, which is approximately ten minutes long (E), can be reduced by incubating ATX with 18:1 LPA for 30 min before addition of LPC (F). Error bars represent the SEM. from three different samples. The data was fitted using the same equation used for LPA inhibition Equation 10.

Taken together, these assays point out to two separate LPA sites: an orthosteric high affinity (^~^50nM) site, virtually identical to the primary LPC binding site that leads to catalysis or competition with minimal inhibitors and has been well characterized ^25,26,34;^ and an allosteric low affinity (^~^1.5 µM) site that leads to an activation of LPC hydrolysis. These observations are of key importance to start understanding the lag phase we described above.

### LPA binds to the ATX tunnel that acts as an allosteric regulatory site

The most likely candidate for an LPA allosteric binding site is the ATX tunnel, which has been shown to bind natural steroids ^16^ and, importantly, has also been suggested to bind LPA in a series of crystal structures reported in ^11^; where the authors showed density for about six carbons of the acyl chain of LPA, bound to the tunnel. Aiming to confirm this interaction, we designed a series of biochemical assays studying the modulation of ATX enzymatic activity by LPA in the presence of different well-characterized inhibitors that bind in the orthosteric and allosteric sites.

The first of these inhibitors was the bile salt TUDCA, an inhibitor with IC_50_ of 11 µM, which we previously reported to bind the allosteric site ^16^. The effect of LPA activation under conditions where ATX is partially inhibited by three TUDCA concentrations was measured. LPA is able to alleviate inhibition by TUDCA, and LPA-mediated activation is more significant at higher concentrations of TUDCA, (Fig.5A-B) suggesting that TUDCA and LPA compete for the same site. In contrast, ATX partially inhibited by PF-8380, a well-established orthosteric site inhibitor ^14,17^ with an IC_50_ of 20 nM, reacts differently to LPA activation (Fig. 5D-E); the activation that LPA is able to exert was roughly the same as in the absence of PF-8380, suggesting no interaction between the two, and that PF-8380 binding does not interfere with LPA-mediated activation. Consistently, when ATX is inhibited by compound 17 ^17^, a high-affinity (11 nM) “hybrid” orthosteric and allosteric site binding inhibitor, LPA activation can no longer be observed, especially at higher compound 17 inhibitor concentrations (Fig.5G-H).

**Figure 5.**
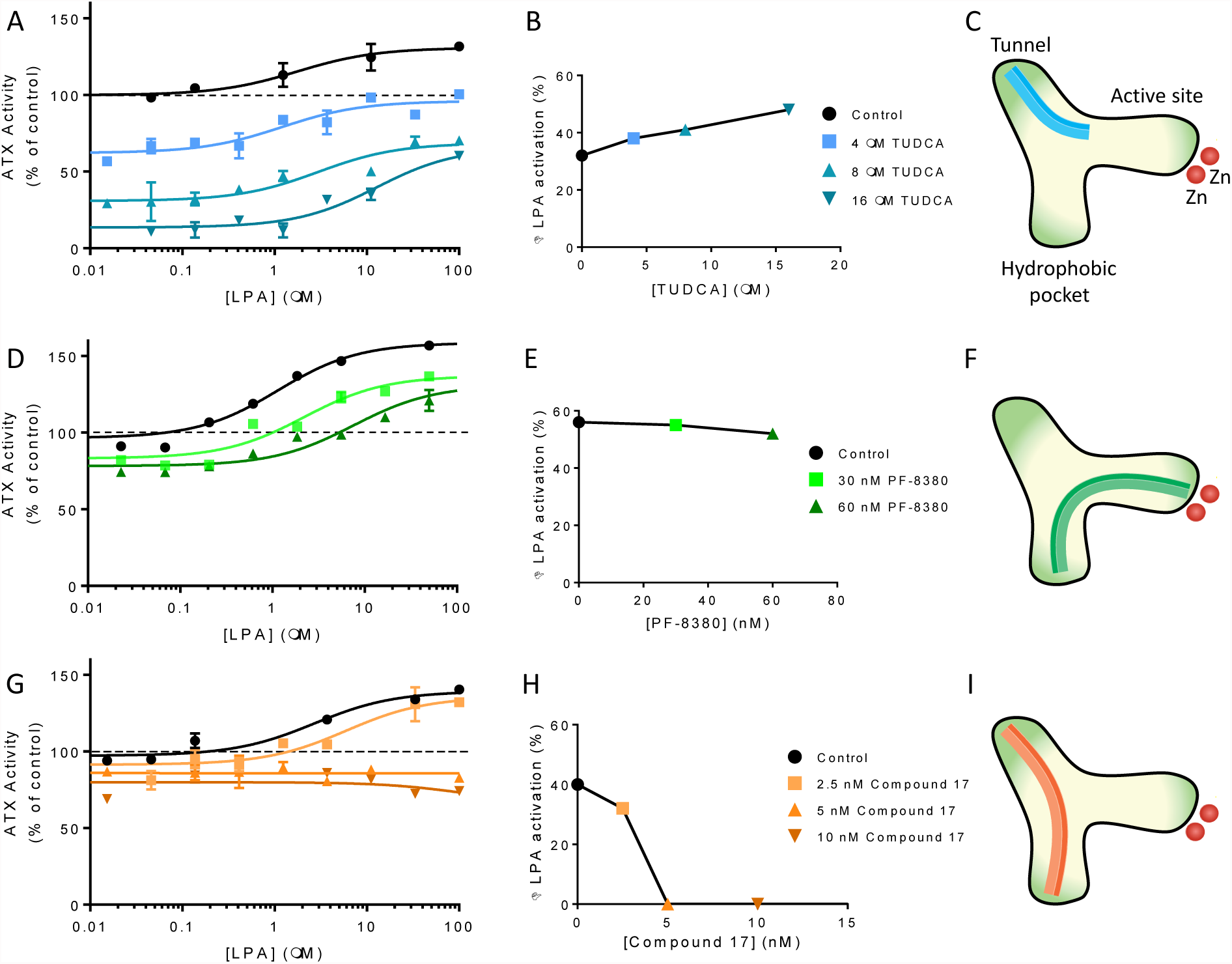
LPA activation in the presence of distinct inhibitors of ATX lysoPLD activity. A, D, G, relative change in activity when LPA was added to the reaction with respect to control activity in the absence of inhibitor. In all cases, 20 nM rat ATX and 150 µM LPC were added to the reaction buffer, and ATX was incubated for 30 min with each inhibitor. Slopes were taken from 60 min after the start of the reaction. B, E, H, increase of ATX activity with respect to the inhibitor-bound ATX control. The data displayed represent the mean value of triplicate measures ± SEM from the fit to Equation 10. C, F, I, schematic representation of the binding sites in ATX structure and the different inhibitor binding modes tested. Type III (allosteric site-binding) (5dlw), I (orthosteric site-binding) (5l0K) and IV (allosteric-orthosteric site hybrid)^17^ inhibitors are represented in cyan, green and orange, respectively. The black dashed lines depict secondary ligands modelled next to the inhibitor.

In summary, we observe that LPA binding to the allosteric site, the tunnel, with an IC_50_ of 1.5 μM, can directly compete with TUDCA binding (IC_50_ = 11 μΜ) and alleviate TUDCA mediated inhibition, but cannot activate PF-8380-mediated ATX inhibition further than the effect it exerts on un-inhibited ATX. Finally, LPA has hardly any effect on ATX inhibited by the strong inhibitor compound 17 (IC_50_ = 16 nM), as it cannot displace it from the orthosteric site binding pocket, nor can it activate by binding to allosteric site tunnel. Collectively, these experiments are all compatible with the hypothesis that LPA binds the tunnel to activate LPC hydrolysis.

### Molecular dynamics simulations support LPA binding to the tunnel and point to its preferred entry route

To further confirm our hypothesis and assess if the tunnel can be occupied by LPA that acts as an allosteric activator, we performed a series of molecular dynamics (MD) simulations. We performed twelve replicas of the same simulation system, which contained the protein, one LPA 18:1 molecule already in the hydrophobic pocket and ten LPA molecules randomly placed in the solvent. In this system, all molecules had different random starting velocities, which were used to produce the MD trajectories. The LPA molecule placed in the hydrophobic pocket remained stable in this location in all simulations (Fig.6A). To illustrate the binding of LPA in the tunnel, we plotted the root mean square distance (RMSD) of all LPA molecules in the solvent from the equivalent atoms of the acyl lipid chain (C10 to C18) residing in the tunnel modelled in PDB entry 3nkp ^11^. In 50% of the simulations, one of the solvent-located LPA molecules moved towards the tunnel within the first 100 ns of the simulation (Fig. 6B). In all binding occurrences, the pathway of entry to the tunnel took place by introducing the hydrophobic tail through the tunnel entry close to the hydrophobic groove of the protein (Supplemental movie 1). Furthermore, once entering the tunnel, lipid molecules remained stable in position through the rest of the simulation, around residues L79, F211, F250, H252, W255, W261 and F275. Interactions of the LPA in the tunnel also involved electrostatic contacts between the phosphate group and R245, R257 and K249 (Fig.6C). Thus, the MD simulations suggest that the preferred entry route of LPA molecules involves a primary sensing of the area of the PDE domain around the tunnel, followed by the introduction of LPA hydrophobic tails into this cavity, to remain bound to ATX. Interestingly, in a 0.5 μs control simulation where the bound LPA was left to equilibrate without additional LPA in the solvent, it remained bound to ATX, and did not leave either by diffusing to the solvent, or moving through the tunnel as suggested in ^11^. The MD simulations support the hypothesis that the low affinity allosteric activation site for LPA is the tunnel.

**Figure 6.**
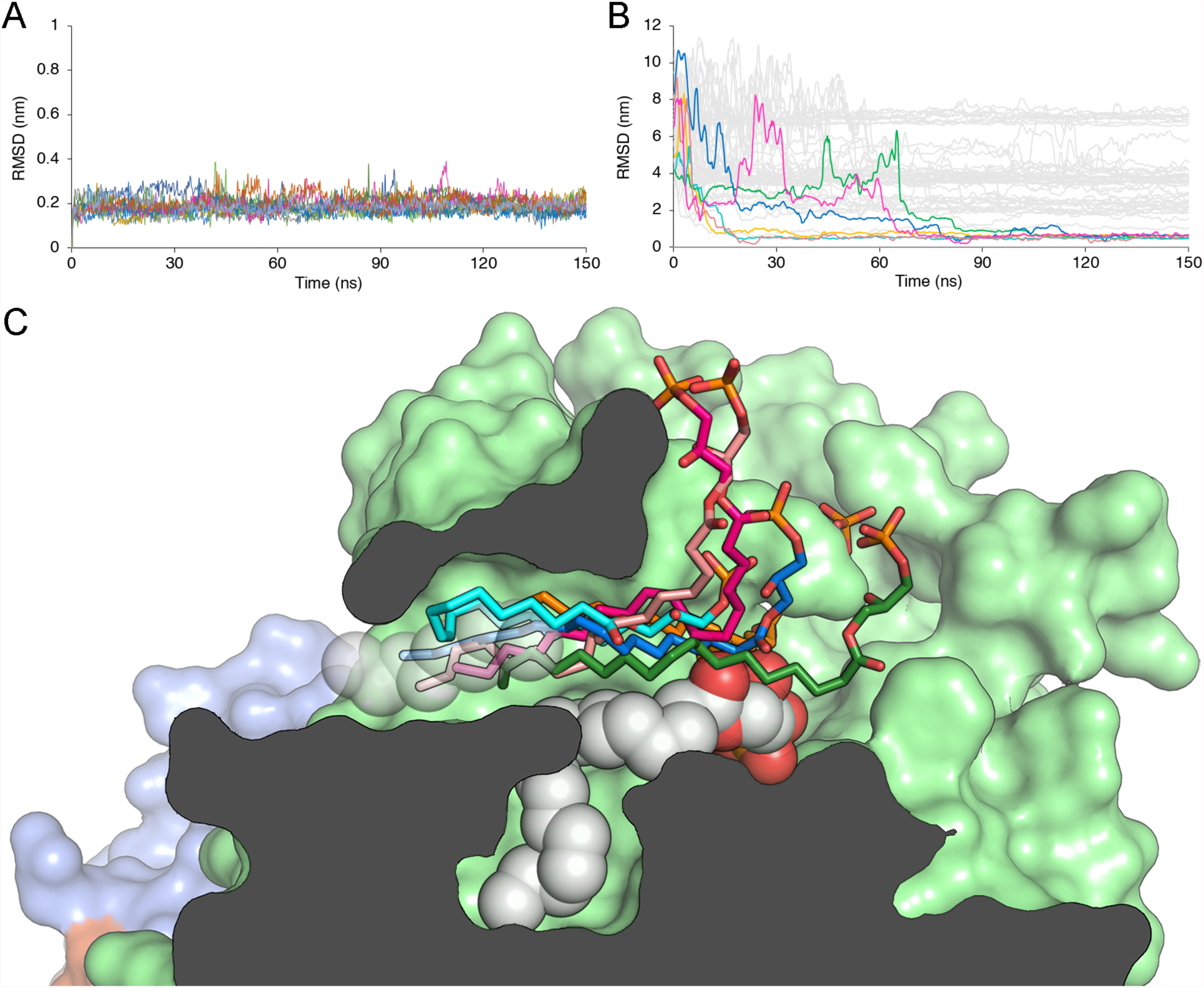
Molecular dynamics simulations of LPA binding to the tunnel of ATX. A, RMSD values of the 18:1 LPA molecule (heavy atoms) bound to the hydrophobic pocket, derived from all simulations after protein backbone fitting. B, RMSD values of the hydrophobic lipid tail (carbon atoms C10-C18) to the reference atoms from the refined 3nkp crystal structure. Coloured lines represent the LPA molecules that entered the tunnel cavity of ATX, corresponding to the colours of Fig.6C, while grey lines represent the remaining LPA molecules (except the one in the hydrophobic pocket) of the system. C, representation of the binding modes of 18:1 LPA molecules to the tunnel. The lipid molecules that entered and remained in the tunnel during the MD simulations are shown in stick representation and colours green, blue, magenta, salmon, cyan and orange. The carbon atoms used for the RMSD calculations are shown in white transparent spheres and the LPA molecule occupying the hydrophobic pocket is shown in white non-transparent spheres representation. The surface of the protein is colour coded as follows, PDE domain in light green, NUC domain in blue and the lasso loop in orange.

### A kinetic model explaining LPA modulation of ATX lysoPLD activity

Having validated our hypothesis that LPA binding to the ATX tunnel leads to activation of the catalytic activity of ATX towards LPC, we employed a bottom-up approach to construct a kinetic model explaining LPA activation. For this, we used the KinTek Global Kinetic Explorer™ software ^35,36^, allowing different types of experimental data to be fitted simultaneously and directly to user-defined reaction models, avoiding simplifying approximations. Our final model, accounting for LPA activation in the ATX catalytic pathway, is expressed in Figure 7.

**Figure 7.**
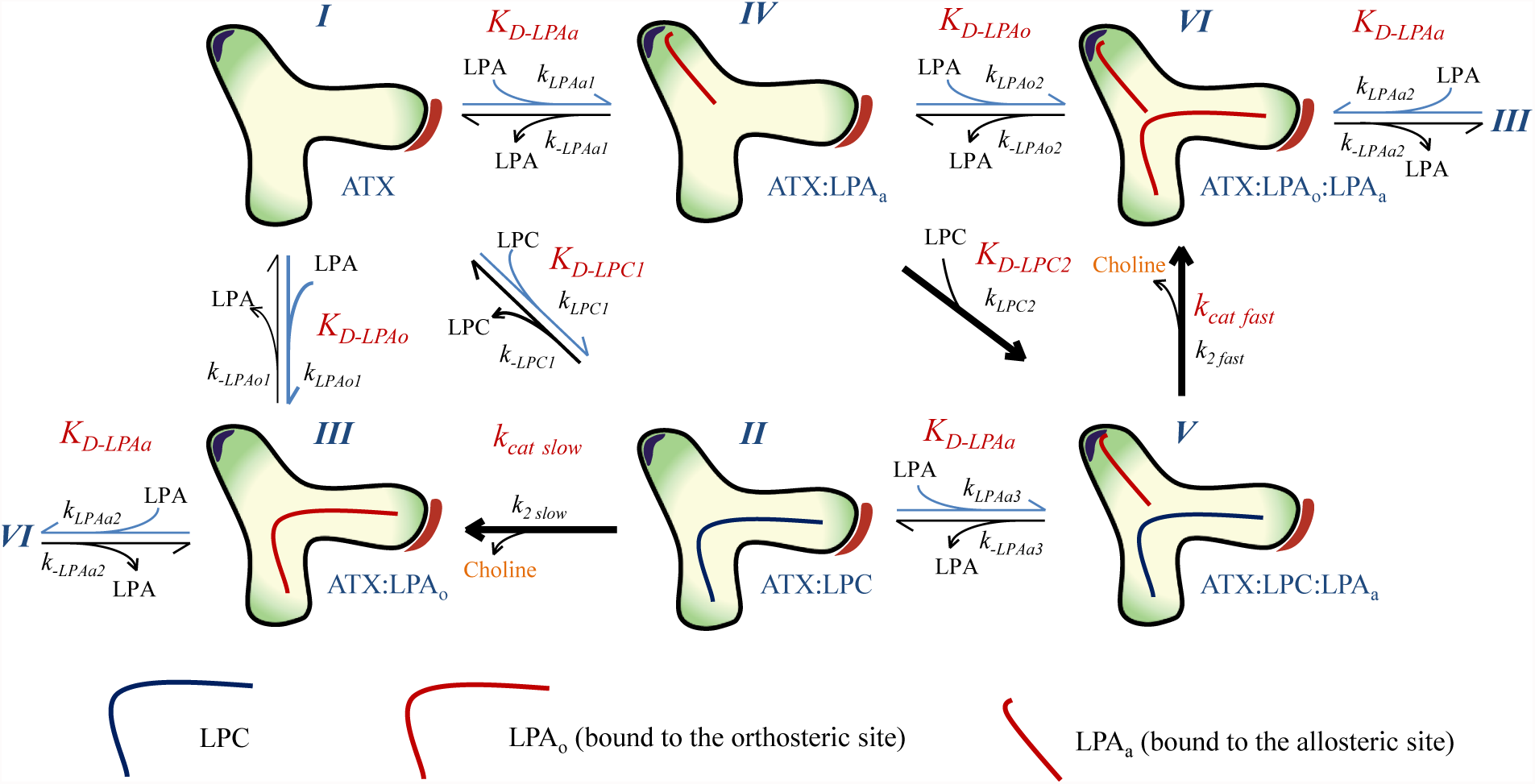
Kinetic model for the modulation of ATX activity by LPA. A, cartoon representing the two ATX kinetic cycles (I-III and IV-VI). Starting from lipid-free ATX [I], LPC binding results to a transient complex [II] that undergoes slow hydrolysis (defined by k_cat slow_): this corresponds to the initial slow phase of in vitro catalysis. Upon catalysis, LPC is converted to choline and LPA, which can remain bound to the orthosteric site [III] or dissociate from ATX, as defined by K_D-LPAo_. This cycle represents the known catalytic pathway for LPC hydrolysis. LPA binding in the tunnel, however, can occur in lipid-free [I], LPC-bound ATX [II] or LPA_o_-bound ATX [III] to yield the respective LPA-carrying species [IV, V & VI]. This binding is defined by K_D-LPAa_, the dissociation constant for LPA in the allosteric site, which occurs independently of the presence of LPC in the orthosteric site. LPC in the presence of LPA bound in the tunnel [V] undergoes faster hydrolysis, defined by k_cat fast_, and yields ATX with two bound LPA molecules [VI]: this is the steady state rate observed in ATX catalysis. Lastly, ATX bound to one LPA molecules in each site (VI) can lose either tunnel-bound LPA as defined by K_D-LPAα_ leading to ΑΤΧ with LPA only bound to the orthosteric site (ΙΙΙ) or it can lose the LPA bound in the orthosteric site as defined by K_D-LPAo_ leading to ΑΤΧ with LPA only bound to the allosteric site (IV).

In addition to reactions directly related to hydrolysis of LPC to LPA and choline by ATX, as choline release is not directly observed, we needed to take into account the rates of the coupled reactions that result to the fluorescent readout of the HVA dimer (HVA^*^). Through a series of titration experiments (for details, see methods), we showed that HVA oxidation by HRP is rapid, and we could describe choline conversion to HVA^*^ with a simplified reaction scheme:

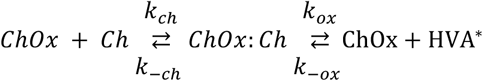

This scheme allowed us to take into account for our model the exact rates of the conversion of choline to the fluorescent signal.

We then collected experimental data of titrations of [LPA], [LPC], and [ATX] versus each other; a complete list of all titrations is available in Methods. In the model, LPA_o_ refers to LPA binding to the orthosteric site (hydrophobic binding pocket) and LPA_a_ refers to LPA binding in the allosteric site (tunnel). Next, having the data and the complete model at hand, we needed to consider simplifications and test assumptions that would allow a robust fit of the most relevant kinetic parameters:

1. All diffusion-limited constants, *k_LPC_, k_LPAο(1-2)_* and *k_LPAa(1-3)_* and *k_ch_* were fixed to 100 µM^-1^ s^-1^
2. The hydrolysis reactions were considered practically non-reversible, and kinetic constants *k_-2slow_*, and *k_-2fast_* were set to zero.
3. The inclusion of the ATX:LPA_o_:LPA_a_ intermediate (VI), the presence of which is supported also by our MD data, is key to explain all data. From that complex, either LPA_o_ or LPA_a_ could leave ATX depending on their relative affinities for the orthosteric or allosteric sites.
4. To better define the binding of LPA to the orthosteric site, we included in the model the data from a titration of LPA inhibiting *p*NP-TMP hydrolysis by competing for binding to the orthosteric site, which allows defining better the affinity of LPA. Further, our model suggests that LPA can be released from the orthosteric site either when it’s the only bound LPA species (III->I), or while another LPA molecule is bound to the allosteric site (VI->IV). To check if the dissociation constants for these two events are different, we first refined them independently; however, they converged to similar values within confidence limits (see Methods). Therefore, these dissociation constants, k_-LPAο1_ and k_-LPAο2_, were constrained to be identical, and we refer to these as k_-LPAo_. In other words, the ability of LPA to bind to the orthosteric site is independent of the presence of LPA in the allosteric site.
5. Then we wanted to check if the ability of LPA to bind to the allosteric site is independent of the presence of LPA in the orthosteric site. LPA can be released from the allosteric site in three instances: when it’s the only bound LPA species (IV->I); when the orthosteric site contains LPA (VI->III); and when the orthosteric site contains LPC (V->II). To check if these three dissociation constants are different, we first refined them independently; however, they converged to similar values within confidence limits (see methods). Therefore, these dissociation constants, k_-LPAa1_, k_-LPAa2_ and k_-LPAa3_ were constrained to be identical and we refer to these as k_-LPAa_. In other words, the ability of LPA to bind to the allosteric site is independent of the presence of LPA or LPC in the orthosteric site.
6. Last, we wanted to check if the ability of LPC to bind to the orthosteric site is affected by the presence of LPA in the allosteric site. When the binding constants for LPC in the absence of LPA in the allosteric site (k_-LPC1_) and in the presence thereof (k_-LPC2_) were released, they did not converge to similar values. Namely, k_-LPC2_ reached zero when fitted, suggesting that LPC does not dissociate when LPA is bound to the allosteric site, and directly implying that the LPA-mediated activation of ATX is also mediated by stabilization of LPC binding.
7. In this model we are not considering the binding of LPC to the allosteric site; this would not be much relevant as it would be unproductive for hydrolysis, and we have no data that would support such a hypothesis. Last but not least such an event is not necessary to explain our data.

Using the FitSpace algorithm included in KinTek Explorer, we analysed the fitted kinetic parameters to obtain their confidence contours (supplemental figure 8). These indicate the extent to which the defined reaction steps and their parameters are constrained by the experimental data ^36^.

Our modelling suggests that LPC can dissociate quickly from the orthosteric site when there is no LPA in the allosteric site, but not when LPA is present in the allosteric site. Thus, binding of LPC to LPA-free ATX can often be unproductive, while binding of LPC to ATX that already carries LPA in the allosteric site, is more likely to be productive. Thus, the transient complex of ATX and LPC alone shows a slow apparent rate of hydrolysis (defined by *k_cot slow_*) corresponding to the initial slow phase of in vitro catalysis, while the complex of ATX, LPC and LPA [V] undergoes faster hydrolysis, as observed in the linear steady state (Figures 2A, 3C, 4EF), and is a consequence of product accumulation as the reaction proceeds. In the absence of any evidence showing a rearrangement of the active site, we conclude that the stabilization of LPC in the orthosteric site by the presence of LPA in the allosteric site is the major reason for the apparent increase in the catalytic rate: the longer residence time of LPC is more likely to lead to be productive.

### Different LPC and LPA species comply with the kinetic model for LPA-mediated activation

As *in vivo* several LPA species co-exist, we next studied whether the product-mediated activation was restricted to the 18:1 species. To that end, the cleavage of five LPC species (14:0, 16:0, 18:0, 18:1 and 22:0) was examined in the presence of their homologous LPA species (14:0, 16:0, 18:0, 18:1 and 22:4). In consequence, AC_50_ values, as well as the % activation were calculated using the Michaelis-Menten derivation (Supplemental figure 4 and supplemental table 4) and modelled using the same procedure and constraints as for LPA 18:1 (Table 3). The results indicated that there is a difference in activation depending on the selected lipid pair, from which the 16:0 and 18:0 species showed a higher activation. Just like 18:1 LPA-mediated activation, the increase in *k_cat_*, as well as no LPC dissociation in the presence of LPA in the allosteric site (i.e. *K_D-LPC2_* converged to zero when fitted), accounted for the activation event of the other species.

**Table 3.** Dissociation constants for LPC and its activating homologous LPA species fitted in the kinetic model (Fig.8).

| LPC/LPA Species | $K_D$-LPC1<br>(μM) | $K_D$-LPA<sub>O</sub><br>(nM) | $K_D$-LPA<sub>A</sub><br>(μM) | $k_{cat}$ slow<br>(s<sup>-1</sup>) | $k_{cat}$ fast<br>(s<sup>-1</sup>) | $k_{cat}$ fast/$k_{cat}$ slow |
| --- | --- | --- | --- | --- | --- | --- |
| 14:0 | 0.54 ± 0.08 | 62 ± 8 | 3.6 ± 0.2 | 0.067 ± 0.021 | 0.10 ± 0.05 | 1.5 |
| 16:0 | 0.7 ± 0.1 | 50 ± 3 | 2.5 ± 0.1 | 0.018 ± 0.011 | 0.21 ± 0.13 | 11.8 |
| 18:0 | 2.36 ± 0.03 | 56 ± 9 | 1.31 ± 0.6 | 0.020 ± 0.009 | 0.18 ± 0.05 | 8.8 |
| 18:1 | 1.71 ± 0.36 | 72 ± 9 | 2.05 ± 1.25 | 0.013 ± 0.003 | 0.043 ± 0.009 | 3.3 |
| 22:0 / 22:4 | 3.08 ± 0.05 | 38 ± 6 | 1.08 ± 0.15 | 0.004 ± 0.018 | 0.008 ± 0.003 | 2.1 |

Next, we examined the activation of the cleavage of all five LPC species available, against all the other LPA species available in a systematic manner. For practical reasons, here we only determined the percentage activation using as normalisation reference the activity of ATX for LPC alone in the start of the reaction. The activation is presented as a 2D heat map plot (Figure 8). The highest activation rates were measured when shorter LPA species were present in the hydrolysis of long LPC species and *vice versa*. This indicates that there is a “cumulative optimal length” of the chain length of the bound LPC and LPA species, likely needed to optimally reinforce the binding of the LPC and aid hydrolysis.

**Figure 8.**
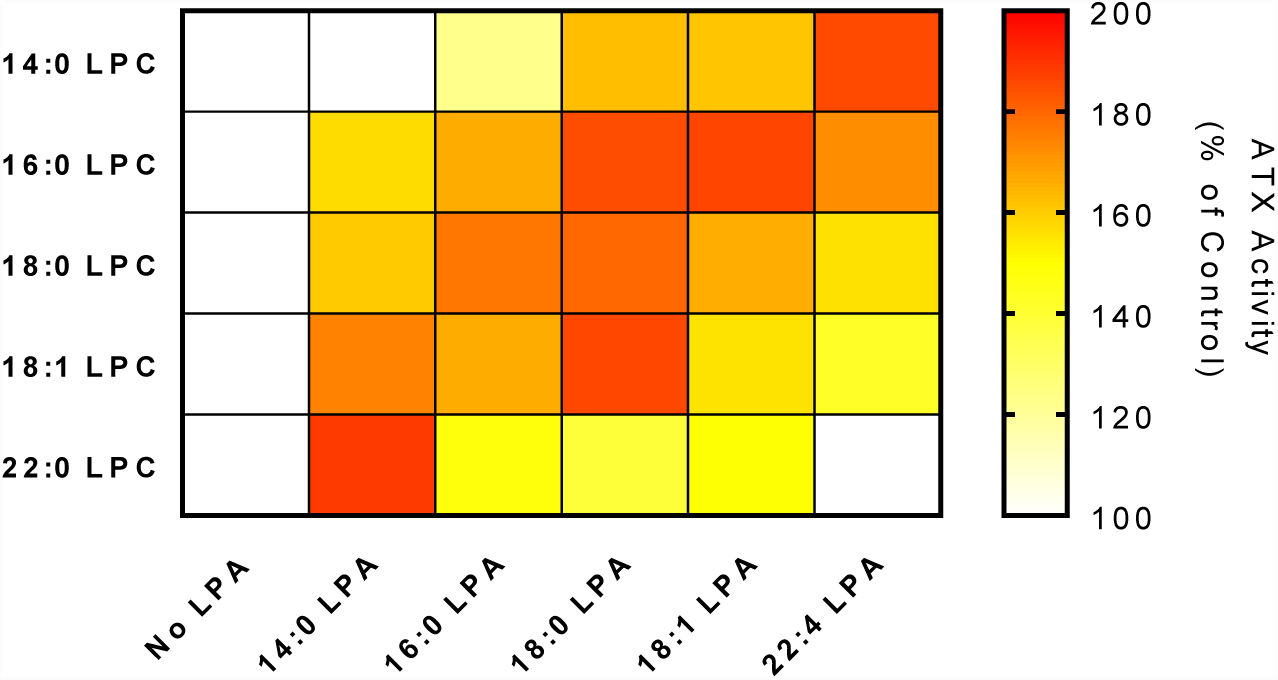
Physiological interplay between shorter and longer-chain LPC and LPA species suggests preferred partnerships in LPA activation. ATX activity changes the slopes measured for the linear phase of ATX lysoPLD activity in the presence of the indicated LPC and LPA molecular species. For each LPC species ATX activity was normalized to the reaction in the absence of LPA, and the maximum increase in activity, as well as the AC50 values (supplemental tables 2 and 3) were calculated form equation 10.

## DISCUSSION

Understanding the mechanisms that underlie the role of LPA signalling in health and disease implies a complete understanding of LPA production of LPC by the ATX lysoPLD activity. While ATX catalysis has been extensively studied in recent years and reaction models have been proposed to explain LPC hydrolysis, many aspects are still poorly understood.

There has been some controversy in the literature, concerning the role of LPA in ATX catalysis. While it is widely accepted that LPA can inhibit hydrolysis of artificial substrates (which we confirm in this study), the role of LPA in LPC hydrolysis has been somewhat controversial. An initial study over a decade ago, showed some LPA product inhibition ^25^, which has later been disputed (**ref**). The data we present here show that the actual mechanism points to the opposing direction: different LPA species are able to activate ATX lysoPLD activity on different natural LPC substrates. On a related note, BrP-LPA has also been shown to inhibit approximately 75% of ATX lysoPLD activity on several LPC molecular species, also affecting plasma ATX activity and LPA levels *in vivo* ^37^. Albeit we note that we have been consistently unable to reproduce that data *in vitro*, we could speculate that the presence of the Br in the head group, might alter the interplay between LPC and LPA, and result in apparent inhibition, at least *in vivo*.

Saunders and colleagues described a model explaining NBD-LPC and FS-3 hydrolysis by ATX ^28^. Specifically, they showed that product release occurs in a sequential manner, where choline is released first and is then followed by LPA release. The discovery of the tunnel as an allosteric site which can modulate ATX activity^16^ offered an opportunity to extend this model. This also made it possible to explain previous observations that we were unable to rationalize before, namely the LPA mediated ATX activation and the lag phase that we have consistently observed in our choline release assays for LPA hydrolysis. The data we present here extend and complement the existing kinetic model, leading to a more complete understanding of LPC catalysis by ATX.

The key element in the new model is indeed the tunnel, an LPA-binding allosteric site. The first suggestion of LPA-binding in the tunnel was reported in the mouse ATX crystal structure by Nishimasu and colleagues ^11^. Our data confirm that LPA binds to the tunnel; However, MD simulations do not support that the tunnel is an exit site for LPA newly formed in the binding pocket, as initially suggested. They suggest that binding of LPA to the tunnel is more likely an independent event. In the absence of data showing that the tunnel can act as an exit site or LPA, the independent binding model should be preferred.

In our new kinetic model, we define two LPA hydrolysis rates, *k_cat fast_* and *k_cat slow_*, which explain the lag phase in ATX kinetics. The faster rate is observed when LPA accumulates (or is added exogenously) and is bound to the tunnel. The observed activation by LPA in the allosteric site can be explained by stabilising the LPC in the orthosteric site, increasing its chances to undergo hydrolysis. This is reflected to our kinetic model, as the affinity of the orthosteric site for LPC is much higher than in the absence of LPA in the allosteric site, suggesting that every single binding even of LPC in presence of LPA is catalytically productive. However, we cannot exclude conformational changes caused by LPA in the allosteric site that decrease the stability of the substrate when bound to the orthosteric site, reducing the activation energy for the catalytic event.

An important aspect of our model is that the activation occurs even between different LPC and LPA molecular species, a condition that reflects the *in vivo* situation. Certain LPC/LPA pairs in the orthosteric and allosteric sites result in a more active ATX. In a physiological situation, the presence of short LPA could accelerate hydrolysis of long LPC, and *vice versa*, creating exciting positive feedback loops between different substrates and products, and perplexing regulatory mechanisms.

Our data lead to two novel hypotheses that should be pursued in ATX research.

First, the allosteric and orthosteric site relationship suggests that different ATX inhibitors, specifically the ones that bind in the tunnel, can affect (positively or negatively) the production of specific LPA species. As it remains unclear how different LPAs signal to the different LPA receptors, such preferential inhibition, implies a likely different clinical outcome for different ATX inhibitors.

Second, it has been shown that ATX binds surface integrins through the SMB domains ^13,31,38^, whereas the longer isoform of ATX (ATXα) binds to heparan sulfate proteoglycans ^32^. While this binding will localize ATX at the cell surface, likely making LPA delivery to surface receptors more efficient, it cannot be excluded that it also affects the kinetics of LPC hydrolysis in our model. Indeed, as the SMB domains are involved in tunnel formation, integrin binding may enable LPA release from the orthosteric or the allosteric site. Taken together, these are exciting research hypotheses that warrant further investigation.

## MATERIALS AND METHODS

### Cells and materials

Human and rat ATX were over-expressed and purified as described in ^39^. Briefly, HEK 293 Flp-In cells (Invitrogen) were grown in Dulbecco’s modified Eagle’s medium (DMEM) containing 10% fetal bovine serum, glutamine and Penicillin-Streptomycin, which were obtained from Thermo Fischer (Massachusetts, USA). Additionally, cell culture was performed in the presence of Hygromycin (Invitrogen). After 4 d in culture, the medium was collected for purification. The culture medium was collected and centrifuged at 4,000 rpm for 15 min. Next, the obtained medium was filtered through a 0.65 µm bottle-top filter, which was subsequently applied at a flow rate of 8–10 ml min^-1^ onto a 10 ml POROS-20 MC column that had been pre-loaded with Ni^2+^. The protein was eluted with 2-3 column volumes of a linear gradient of imidazole. Next, fractions were applied onto a Superose 6 10/30 size-exclusion column and concentrated afterwards.

For the kinetic studies, LPC (18:1), LPA (14:0, 16:0, 18:0, 18:1), 12:0 lyso-NBD-PC, 18:1 NBD-LPE and 18:1 NBD-PEG4 LPE were obtained from Avanti Polar Lipids Inc. (Alabaster, AL). Choline oxidase, Horseradish peroxidase, Homovanillic acid, Sodium TUDCA, choline chloride and fatty acid-free BSA were purchased from Sigma-Aldrich (St. Louis, Missouri). Hybrids and small-molecule ATX inhibitors were synthesized by the group of Craig Jamieson at the University of Strathclyde. Steady-state experiments were performed in Corning^®^ 96- or 384-well OptiPlate from Sigma-Aldrich.

### Kinetic measurements of ATX lysoPLD activity

The biochemical studies of ATX lysoPLD activity were performed with rat ATX (863 aa). Activity was measured by a coupled reaction with 1 U ml^-1^ choline oxidase and 2U ml^-1^ horseradish peroxidase (HRP) and 2 mM homovanillic acid (HVA). For the assays, 18:1 LPC was incubated with 20 nM ATX (obtained from HEK 293 Flip-In cells), reaching a final volume of 100 µl buffer, which contained 50 mM Tris, 0.01%, 50 mM CaCl2, Triton X-100, pH 7.4. Steady-state choline release was measured at 37 °C by HVA fluorescence at λ_ex_ / λ_em_ = 320/460 nm every 30 s for at least 60 min with a Pherastar plate reader (BMG Labtech) ^16^. Due to the presence of a lag phase during the first 10 min of ATX-dependent LPC hydrolysis (Fig.1B), the subsequent linear slope was used to perform all analyses.

For comparison between double exponential 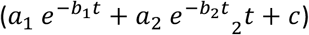 and single exponential 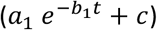 fluorescence kinetics over time, Akaike’s Informative Criteria were used to establish the significance of the fits.

The resulting fluorescence intensity was converted to choline concentration by using a standard curve. This was prepared by titrating increasing concentrations of choline chloride in the reaction buffer in the absence of both LPC and ATX and measuring the end point of its conversion to the chromophoric form of HVA. The data were plotted with PRISM version 4 or 5 (GraphPad Software, San Diego, CA, USA) and presented as mean ± standard error of the mean (SEM).

The slopes of ATX hydrolysis of 18:1 LPC, measured in choline concentration over time, were plotted against substrate concentration to obtain the kinetic parameters by means of nonlinear regression to Michaelis-Menten equation:

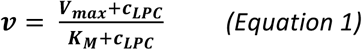

where, *v* represents velocity, measured in choline release over time, *V_max_* is the maximum velocity obtained at a specific enzyme concentration, and *K_M_* is the concentration of substrate (c_LPC_) at which half of *V_max_* is reached.

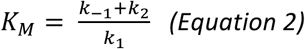

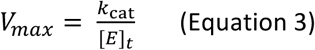

### Measure of NBD-PEG4 LPE steady-state cleavage

Assays to assess ATX lysoPLD activity to hydrolyse the novel substrate NBD-PEG4 LPE were performed in the same conditions as described for LPC hydrolysis. Activity was monitored at λ_exc_ / λ_em_ = 468/546 nm. However, because of solubility problems, NBD-PEG4 LPE was dissolved in Ethanol:H_2_O (1:1) 0.01% TX100. The slopes of the linear part of ATX activity were analysed using Graphpad Prism software to determine inhibition or activation in the distinct assays.

### Inhibition of ATX lysoPLD activity

In order to determine the IC50 for the different inhibitors on lysoPLD ATX activity in the choline oxidase coupled assay, the velocity of the reaction was monitored for each compound as a function of time and the linear phase of the kinetics was taken from 60 min after the addition of ATX to the reaction buffer. The resulting fluorescence intensity signal over time was used to model all inhibitor concentrations simultaneously using the following formula ^40^:

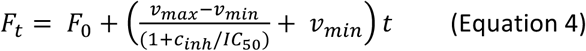

where F_t_ represents the observed fluorescence signal at time (t), F_0_ is the background fluorescence signal at the start of each measurement, v_max_ and v_min_ were fitted for the minimum and maximum relative velocity, and c_inh_ corresponds to the inhibitor concentration for each assay. This equation allowed the calculation of the IC_50_ of each inhibitor. The previous equation can be derived by linear regression of each inhibitor concentration:

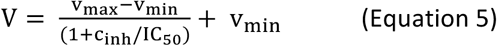

### Mechanistic Studies with ATX inhibitors

For initial comparison between competitive, uncompetitive and non-competitive inhibition, assays measuring LPC hydrolysis by ATX lysoPLD activity were performed and analysed by nonlinear regression. For this purpose, three inhibitor concentrations, determined from the experimentally calculated IC_50_, and an additional control in the absence of inhibitor were used to obtain the slopes from the linear phase of ATX kinetics, as explained above. Next, the following equations were employed to describe each mode of inhibition ^16^:

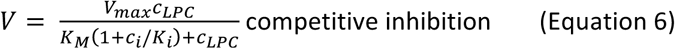

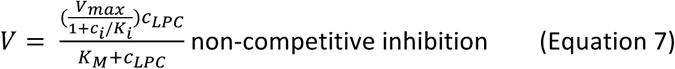

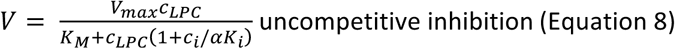

where V is the observed velocity and c_LPC_ is the corresponding LPC concentration for each data point, and c_i_ is the inhibitor concentration for each curve; and *K_i_* is the inhibition constant. Note that equation denotes *K_i_* for competitive inhibition (i.e. binding in the absence of the substrate) and for uncompetitive inhibition (i.e. binding only in the presence of the substrate), where *α* will determine the type of inhibition: *α* < 1 for uncompetitive inhibition, *α* > 1 for competitive inhibition, and *α* = 1 for noncompetitive inhibition.

To calculate the percentage of certainty for each type of inhibition, the alpha value was calculated in the partial mixed inhibition model (Equation 9), and the significance of the analysis was assessed by Akaike’s Informative Criteria.

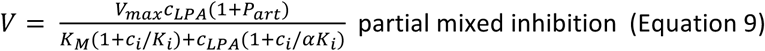

where the new parameter *Part* was defined as the partiality of the inhibition.

### LPA Activation measurements

The activation assays using LPA were performed in a similar way as those done for the inhibitors. In this case, LPA was dissolved in Ethanol:H_2_0 (1:1), 0.01% TX-100 and was added to the reaction buffer. The presence of ethanol was taken into account and controls in the absence of ATX and/or LPC were employed to correct the kinetic data. Normally, the slopes were obtained from 60 min after ATX was added to the reaction buffer and were related to ATX activity in the same conditions but in the absence of LPA. In order to assess whether LPA activation could also take place with different LPA species, 14:0, 16:0, 18:0, and 18:1 LPA were diluted threefold, and data was analysed as described before.

### Mechanistic study of LPA activation

To discriminate between the different structural components that take part in LPA activation of ATX lysoPLD activity, ATX was incubated for 30 min with different concentrations of inhibitors. ATX was subsequently added to the reaction buffer containing 150 µM 18:1 LPC, and 18:1 LPA was diluted threefold. The slopes were calculated from at least 60 min after the reaction was started. The calculation of the percentage of activation by LPA was related to ATX in the same conditions but in the absence of LPA and inhibitors, which represented 100% activity. AC_50_ was obtained from the following equation:

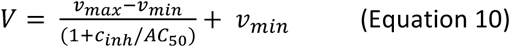

### Modelling on KinTek Explorer

KinTek Global Kinetic Explorer™ (version 6.3) was used to design reaction models, from which simulations could be performed. ATX catalysis models were written and tested in this software, following the workflow described in Supplemental figure 1. The specific steps that were taken in each stage of the design is specified in the results section. Experimental data was fitted directly to the reaction model upon numerical integration of the rate equations using SpectraFit™. Additionally, the robustness of the described model was statistically analysed using the FitSpace Explorer™ software. This was used to define the confidence contour analysis, which allowed establishing the Chi^2^ threshold at the boundaries between the different kinetic parameters. Consequently, complex relations between the kinetic constants can be detected and the extent to which the parameters are constrained by the experimental data can be addressed.

Two titration experiments, namely [Choline oxidase] vs. [Choline chloride] and [HRP] vs. [choline chloride], were performed to simplify the coupled reaction to detect choline release.

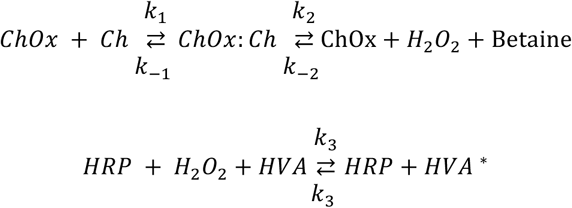

For the modelling of *K_D-LPAo_*, we performed titrations of [LPA] vs. [*p*NP-TMP]. Next, *K_D-LPC1_* and *K_D-LPC2_* were modelled by the titration of [LPC] vs. [ATX]. Lastly, LPA activation was explained by the titrations of [LPA] vs. [ATX], and [LPA] vs. [LPC], which allowed defining *k_catslow_*, *k_catfast_*, as well as *K_D-LPAa_*.

### Molecular dynamics simulations

A structural model of human autotaxin (Uniprot code Q13822) in complex with 18:1 LPA was constructed with Modeller v9.7 using our previously reported crystal structures of rat autotaxin (PDB codes: 5dlw and 5dlt) as templates (94% sequence identity) ^16,41^. All molecular dynamics (MD) simulations were performed using the GROMACS software v5.1 ^42^. The ATX-LPA complex was then inserted in a pre-equilibrated box containing water. Apart from the 18:1 LPA molecule in the hydrophobic site, ten extra 18:1 LPA molecules were added to the simulation box in random positions, with a minimum distance of 40 Å to the tunnel. The AMBER99SB-ILDN force field was used for all the MD simulations along with the TIP3P water model ^43^. Force field parameters for the lipid molecules were generated using the general Amber force field (GAFF) and HF/6-31G*-derived RESP atomic charges ^44^. The reference system consisted of the protein, eleven 18:1 LPA molecules, ~58.500 water molecules and 17 Na^+^ ions (for total charge equilibrium) in a 12.5 × 12.5 × 12.5 nm simulation box, resulting in a total number of 189032 atoms. The system was energy minimized and subsequently subjected to a 10 ns MD equilibration, with an isothermal-isobaric ensemble, using isotropic pressure control and positional restraints on protein and lipid molecule coordinates. The resulting equilibrated system was replicated 12 times, random initial velocities were produced for each one of the replicas using a random seed and independent 150 ns MD trajectories were produced in a constant temperature of 310K, using separate v-rescale thermostats for the protein, the lipids and solvent molecules. A time step of 2 fs was used and all bonds were constrained using the LINCS algorithm. Lennard-Jones interactions were computed using a cut-off of 10 Å, and the electrostatic interactions were treated using PME with the same real-space cut-off. A control 500 ns simulation was also run without the ten surrounding LPA molecules, to examine the stability of LPA in the orthosteric pocket. All runs added up to a total simulation time of 2.4 μs.

## Acknowledgements

We thank Wouter Moolenaar for his continued support and enthusiasm and for critically reading this manuscript. An early part of this work was supported by an NWO-TOP grant to AP (700.10.354). We would like to thank Prof. Leonardo Pardo for providing infrastructure in the Laboratory of Computational Medicine, UAB, for the molecular dynamics simulations.

